# Astrocyte CB1 receptors are required for inhibitory maturation and ocular dominance plasticity in the mouse visual cortex

**DOI:** 10.1101/2023.12.10.570998

**Authors:** Rogier Min, Yi Qin, Sven Kerst, M. Hadi Saiepour, Mariska van Lier, Christiaan N. Levelt

## Abstract

**Summary and Introduction:** Neuronal circuits are shaped by experience. This happens much more readily in the young compared to the adult brain. The unique learning capacity of the young brain is regulated through postnatal critical periods, during which the ability of neuronal networks to re-wire is greatly enhanced^1^. Endocannabinoids, which signal through the cannabinoid CB1 receptor (CB1R), regulate several forms of neuronal plasticity^2^. In the developing neocortex, CB1Rs play a key role in the maturation of inhibitory circuits. For example, interfering with CB1R signaling during development disrupts inhibitory maturation in the prefrontal cortex^3^. In developing primary visual cortex (V1), endocannabinoid-mediated plasticity at inhibitory synapses regulates the maturation of inhibitory synaptic transmission, shifting synapses from an immature state characterized by strong short-term depression to a mature state with reduced short-term depression^4,5^. This maturation step correlates with the timing of the critical period. While CB1Rs were originally thought to reside mainly on presynaptic axon terminals, recent studies have highlighted an unexpected role for astrocytic CB1Rs in endocannabinoid mediated plasticity^6-8^. Here, we investigate the impact of cell-type specific removal of CB1Rs from interneurons or astrocytes on development of inhibitory synapses and network plasticity of V1. We show that removing CB1Rs from astrocytes interferes with maturation of inhibitory synaptic transmission in V1. In addition, it strongly reduces ocular dominance (OD) plasticity during the critical period. In contrast, removing interneuron CB1Rs leaves these processes intact. Our results reveal an unexpected role of astrocytic CB1Rs in critical period plasticity in V1, and highlight the involvement of glial cells in the plasticity and synaptic maturation of sensory circuits.

## Results and discussion

### Removal of astrocyte vs interneuron CB1 receptors

Inhibitory synapses in primary visual cortex (V1) undergo a CB1 receptor (CB1R)-dependent maturation step during postnatal development: they shift from an immature state characterized by strong short-term synaptic depression towards a state with reduced short-term depression^4,5^. This process is complete around P28, corresponding to the peak of the critical period for OD plasticity in mouse^9^. Inhibitory maturation is absent in full CB1R knockout mice^4^. CB1Rs are traditionally thought to reside on presynaptic axon terminals. However, the parvalbumin-expressing fast-spiking interneurons that undergo developmental maturation^5,10^ are thought to express no or only low levels of CB1Rs^11-13^. Furthermore, it was previously shown that astrocytes also express CB1Rs^6,7^, which are involved in plasticity of developing sensory circuits^7^. To investigate how removal of CB1Rs from different cell types (astrocytes vs interneurons) affects inhibitory maturation in V1, we made use of conditional knockout mice lacking CB1Rs in either astrocytes or interneurons. We crossed transgenic mice containing a floxed CB1R gene (*Cnr1*^flox/flox^ mice)^14^ with different Cre-driver lines. For interneuron-specific recombination we used GAD2-Cre knock-in mice^15^, while for astrocyte specific recombination we used GLAST-CreERT2 transgenic mice (JAX stock #012586). The resulting mice lacked CB1Rs either in astrocytes (“GLAST-CB1R-KO mice”) or interneurons (“GAD2-CB1R-KO mice”).

Conditional recombination in GLAST-CreERT2 mice requires induction by tamoxifen injection. For our experiments, astrocyte-specific recombination needed to be induced at a young age, before the start of the critical period. A potential problem with early tamoxifen injection may be that recombination occurs in neuronal precursor cells, leading to recombination in neurons. We therefore tested at which age GLAST-CreERT2 induction was specific for glial cell types. Using a TdTomato Cre-reporter line crossed to GLAST-CreERT2 mice, we found that a single intraperitoneal (i.p.) tamoxifen injection at P1 indeed resulted in recombination in a small number of neocortical neurons (Fig. 1A). In contrast, a single injection between P3 and P5 resulted in efficient and specific recombination in glial cells, with no neuronal recombination in V1. Of all V1 astrocytes 80% showed recombination, while 77% of recombined cells were astrocytes (Fig. 1B,E), the rest being other glial cells (oligodendrocytes, oligodendrocyte precursor cells and NG2 cells; Fig. 1C,D). Because astrocytes are the main glial cell type expressing CB1Rs (Allen Brain Atlas; www.brain-map.org), phenotypic changes observed in GLAST-CB1R-KO mice that are treated with tamoxifen at P3-5 are most likely due to loss of CB1R expression in astrocytes.

**Figure 1.**
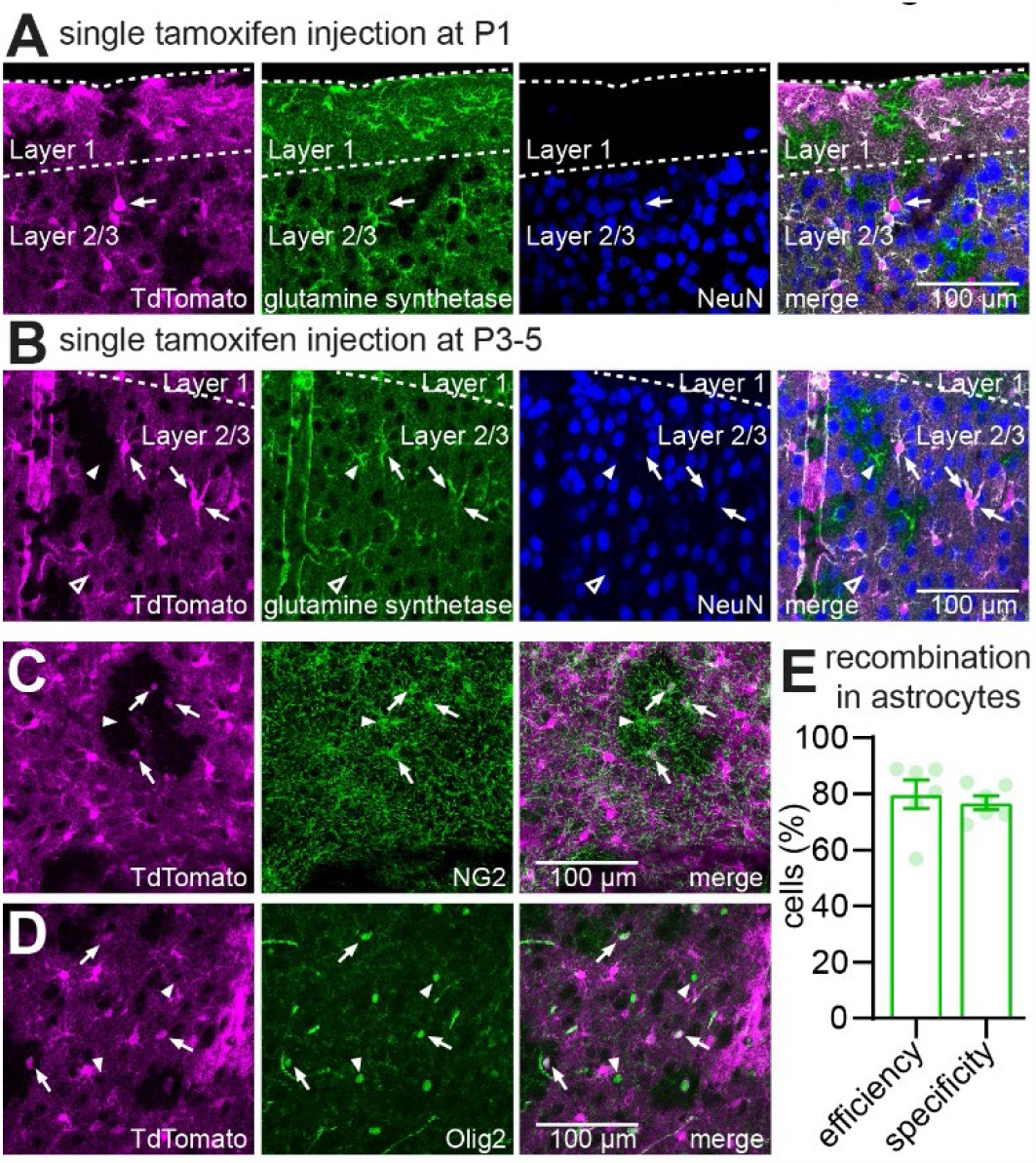
Early astrocytic recombination in GLAST-CreERT2 mice. (A) GLAST-CreERT2 TdTomato mice received a single i.p. injection of tamoxifen at P1. Slices containing V1 were prepared at P28-35, processed for immunofluorescence imaging and visualized using confocal microscopy. Recombination (indicated by TdTomato expression, magenta) was observed in astrocytes (visualized using a glutamine synthetase antibody, green), but also in sparse neurons (visualized using NeuN antibody, blue). An example neuron with pyramidal morphology is indicated by the arrow. (B) Changing the tamoxifen injection regime to a single injection at P3-5 abolished neuronal recombination in V1, while astrocyte recombination was efficient (∼80%). Arrows indicate TdTomato expressing astrocytes, arrowheads indicate TdTomato negative astrocytes. (C,D) Specificity of recombination was high for astrocytes, but some non-neuronal recombination was seen in glial cells positive for NG2 positive cells (C, green, arrows) or Olig2 positive cells (D, green, arrows). Arrowheads indicate TdTomato negative NG2 and Olig2 positive cells. (E) Quantification of efficiency and specificity of recombination in astrocytes in mice receiving a single tamoxifen i.p. injection at P3-5, based on TdTomato and glutamine synthetase positivity.

### Loss of astrocytic CB1Rs interferes with inhibitory synaptic maturation

To investigate how loss of CB1Rs from specific cell types affected inhibitory synaptic maturation we assessed short-term dynamics of inhibitory synapses in acute brain slices of P28-35 mice. Whole cell patch-clamp recordings were made from L2/3 pyramidal neurons, and evoked inhibitory postsynaptic currents (IPSCs) were measured upon repetitive extracellular stimulation (10 pulses at 25 Hz; see methods for recording details). V1 inhibitory synapses onto L2/3 pyramidal neurons normally mature towards a state characterized by less pronounced short-term synaptic depression at P28-35^4,5^. This inhibitory maturation is absent in full CB1R knockout mice, with inhibitory synapses maintaining an immature state characterized by stronger short-term depression^4^. We found that short-term dynamics of inhibitory synapses in P28-35 GAD2-CB1R-KO mice did not differ from that in wild-type littermates (Fig. 2A; normalized steady state IPSC amplitude: wild-type: 0.42±0.05, n=16; GAD2-CB1R-KO: 0.45±0.05, n=19; *P*=0.71; Mann-Whitney test), suggesting normal inhibitory maturation in the absence of interneuron CB1Rs. In contrast, GLAST-CB1R-KO mice showed more pronounced short-term depression when compared to wild-type littermates (Fig. 2B; normalized steady state IPSC amplitude: wild-type: 0.48±0.03, n=11; GAD2-CB1R-KO: 0.33±0.03, n=14; *P*=0.008; Mann-Whitney test). This suggests that loss of CB1Rs on astrocytes, but not on interneurons, interferes with the maturation of inhibitory synaptic transmission.

**Figure 2.**
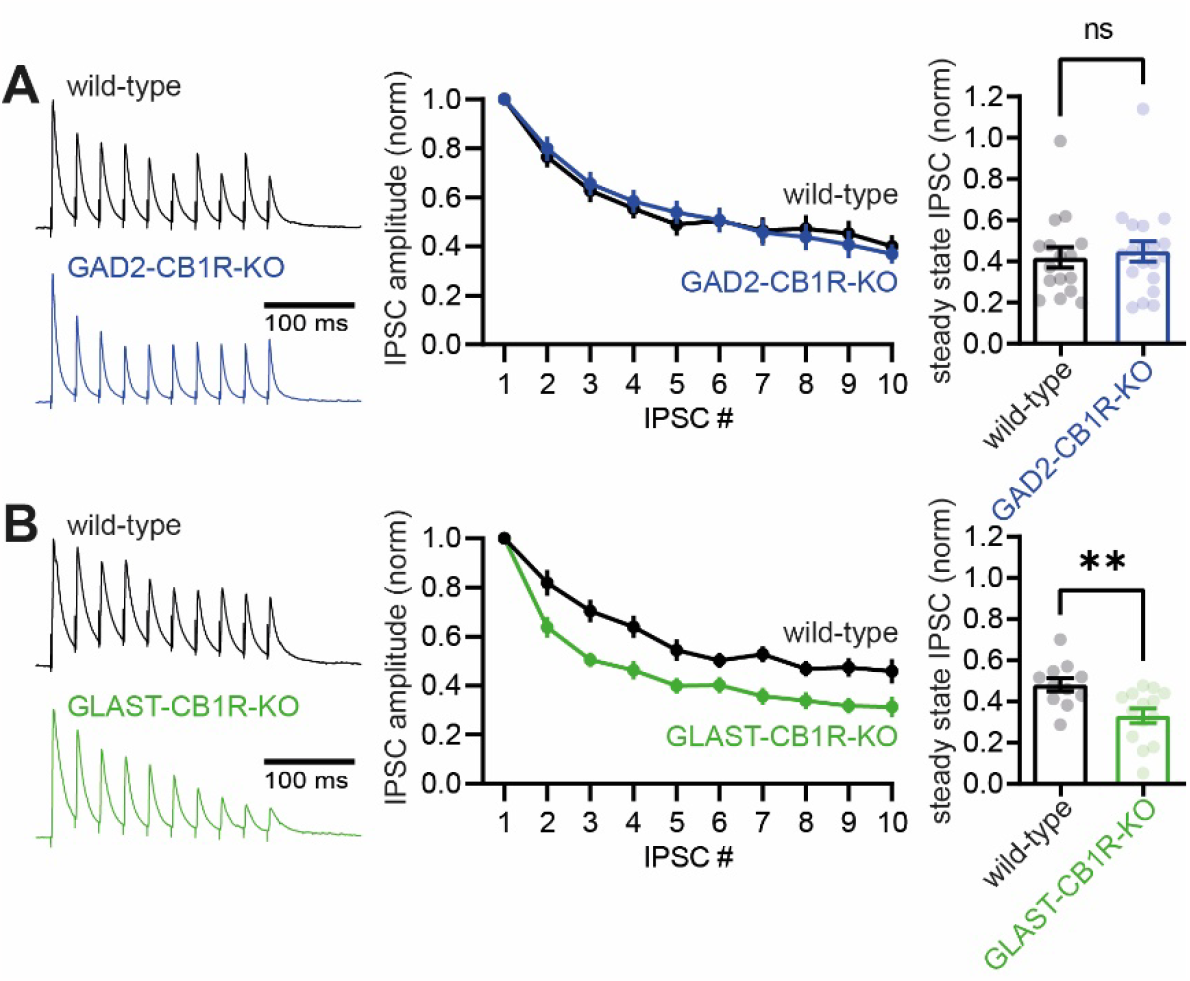
Impaired inhibitory synaptic maturation upon loss of astrocyte CB1 receptors. (A) Left: example traces showing the dynamics of inhibitory synaptic transmission in acute brain slices from GAD2-CB1R-KO mice (blue) and their wild-type littermates (black). Middle: averaged IPSC amplitude normalized to the first, for each of the 10 IPSCs in the train. Right: Steady state IPSC amplitude (averaged normalized amplitude of the last three IPSCs in the train) for all individual recorded neurons (dots). Bars show mean. (B) Same as in A, but for GLAST-CB1R-KO mice (green) and their wild-type littermates (black). Error bars indicate SEM.

### Long-term depression of inhibitory synapses is intact upon removal of interneuron or astrocyte CB1Rs

Inhibitory synapses in V1 can undergo endocannabinoid-mediated long-term depression (iLTD) at early developmental stages, but this form of plasticity is lost during maturation. iLTD is blocked by CB1R antagonists, and absent in full CB1R knockout mice^4,5^. The significance of iLTD for inhibitory circuit functioning and maturation is unknown. To investigate how iLTD was effected by cell-type specific CB1R removal we prepared acute brain slices from young mice (P14-21), and performed whole cell patch-clamp recordings from L2/3 pyramidal neurons. Evoked IPSCs were recorded for a baseline period of 10 minutes, followed by iLTD induction using a theta-burst protocol (see methods for additional details). In wild-type mice this led to a significant reduction in IPSC amplitude, indicating robust iLTD expression (iLTD: 17.0±2.9%, n=29, IPSC amplitude baseline vs after iLTD induction: *P*=0.0002, paired t-test; Fig. 3A). iLTD was abolished in the presence of the CB1R antagonist AM251 (10 μM; iLTD: 0.0±4.2% iLTD, n=12, IPSC amplitude baseline vs after iLTD induction: *P*=0.92; % iLTD control vs +AM251: *P*=0.002, unpaired t-test; Fig 3A).

**Figure 3.**
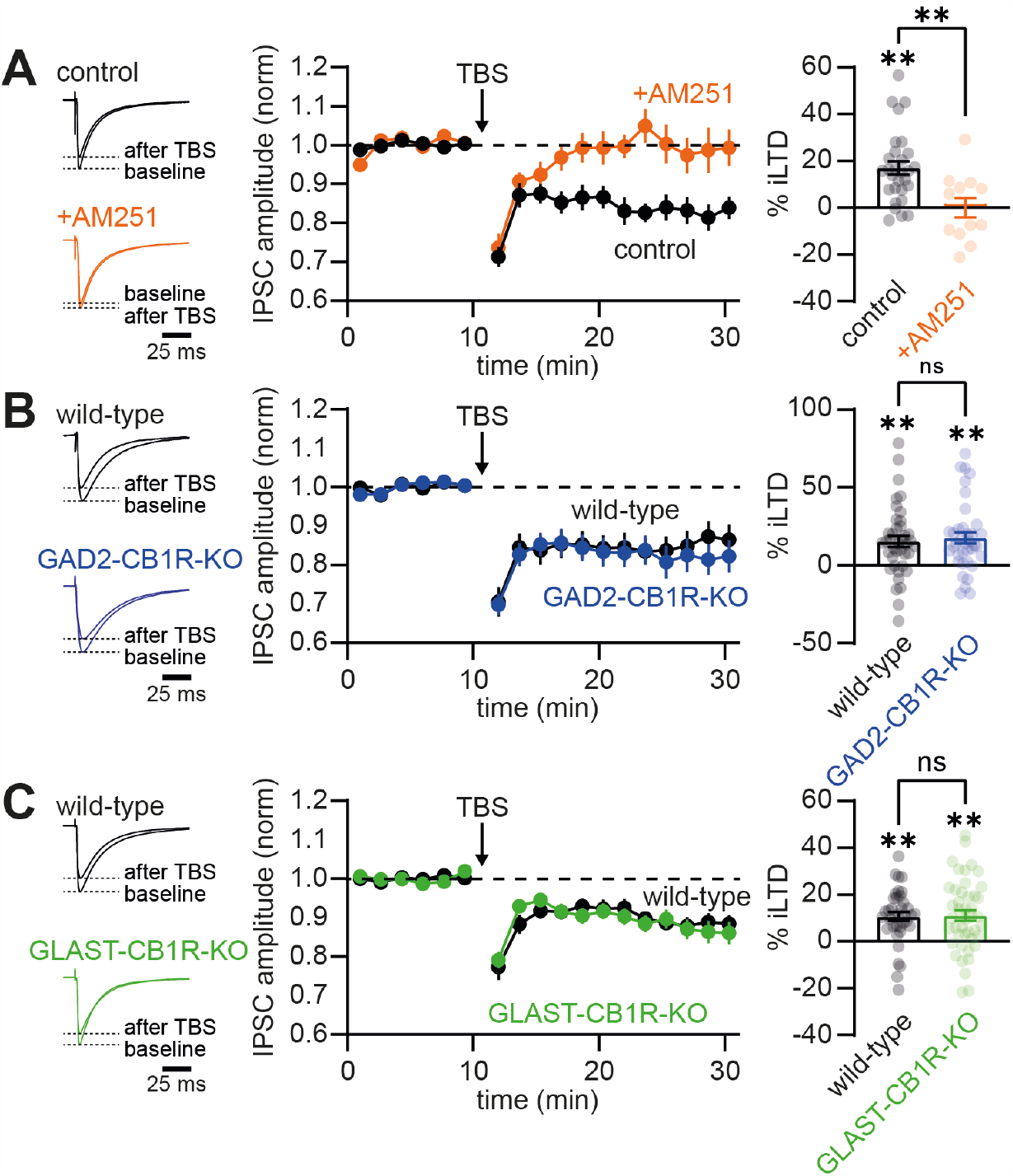
iLTD is unaffected by removal of astrocyte or interneuron CB1 receptors. (A) Left: example traces showing the averaged IPSC during the 10 minutes of baseline recording and 10-20 minutes after iLTD induction by TBS. Black traces are from a control experiment, orange in the presence of the CB1 receptor antagonist AM251. Middle: Averaged time course of the IPSC amplitude normalized to baseline for all experiments under control conditions (black) and in the presence of AM251 (orange). Right: Averaged amount of iLTD (% reduction of the IPSC amplitude after TBS) for all individual recorded neurons (dots). Bars show mean. (B) Same as in A, but now for GAD2-CB1R-KO mice (blue) and their wild-type littermates (black). (C) Same as in A and B, but now for GLAST-CB1R-KO mice (green) and their wild-type littermates (black). Error bars indicate SEM.

Next, we investigated how iLTD was influenced by cell-type specific removal of CB1Rs. Surprisingly, we found that neither removal of interneuron CB1Rs, nor of astrocyte CB1Rs, affected the magnitude of iLTD (Fig. 3B,C). iLTD did not significantly differ between interneuron CB1R knockouts and wild-type littermates (GAD2-CB1R-KO iLTD: 17.7±3.7%, n=37, IPSC amplitude baseline vs after iLTD induction: *P*=0.0002, paired t-test; wild-type littermates iLTD: 15.3±3.6% iLTD, n=40, IPSC amplitude baseline vs after iLTD induction: *P*<0.0001; % iLTD wild-type vs GAD2-CB1R-KO: *P*=0.65, unpaired t-test; Fig. 3B). The same was true for astrocyte CB1R knockouts (GLAST-CB1R-KO iLTD: 11.1±2.3%, n=44, IPSC amplitude baseline vs after iLTD induction: *P*<0.0001, paired t-test; wild-type littermates iLTD: 10.7±1.9% iLTD, n=37, IPSC amplitude baseline vs after iLTD induction: *P*<0.0001; % iLTD wild-type vs GLAST-CB1R-KO: *P*=0.91, unpaired t-test; Fig. 3C). Therefore, while inhibitory synaptic maturation relies on astrocyte CB1Rs, iLTD is surprisingly intact upon conditional removal of either astrocyte or interneuron CB1Rs.

### Loss of astrocytic CB1Rs disrupts OD plasticity

The maturation of inhibitory synaptic transmission is known to be critical for the occurrence of OD plasticity during the critical period^1^. Therefore, we assessed OD plasticity in mice in which CB1R expression was disrupted in astrocytes or interneurons. Using optical imaging of intrinsic signal^16^, we measured responses to stimulation of the two eyes in the binocular region of V1, calculated the OD index (ODI; see methods), and compared mice that were reared normally (non-deprived) with mice that were monocularly deprived for three days starting around P28 (3 days MD). In wild-type littermates of both GAD2-CB1R-KO and GLAST-CB1R mice 3 days MD led to an OD shift (GAD2-CBR1-KO littermates: ODI non-deprived: 0.34±0.04, n=5; 3 days MD: 0.07±0.05, n=6; GLAST-CB1R littermates: ODI non-deprived: 0.33±0.04, n=6; 3 days MD 0.02±0.06, n=7; Fig. 4A,B). This OD shift was also observed upon interneuron specific CB1R removal (GAD2-CB1R-KO: ODI non-deprived: 0.31±0.04, n=5; 3 days MD: 0.03±0.05, n=5; Fig. 4A). Statistical analysis yielded no interaction between genotype and molecular deprivation for GAD2-CB1R-KO mice (two-way ANOVA; interaction of genotype with OD shift *P*=0.93; Tukey’s post-hoc test: wild-type non-deprived vs 3 days MD: *P*=0.002; GAD2-CB1R-KO non-deprived vs 3 days MD: *P*=0.003; wild-type 3 days MD vs GAD2-CB1R-KO 3 days MD: *P*=0.95). In contrast, no significant OD shift was observed upon removal of astrocyte CB1Rs (GLAST-CB1R-KO: ODI non-deprived: 0.31±0.04, n=7; 3 days MD: 0.22±0.04, n=8; two-way ANOVA; interaction of genotype with OD shift *P*=0.022; Tukey’s post-hoc test: wild-type non-deprived vs 3 days MD: *P*=0.0006; GLAST-CB1R-KO non-deprived vs 3 days MD: *P*=0.54; wild-type 3 days MD vs GLAST-CB1R-KO 3 days MD: *P*=0.021; Fig. 4B). Therefore, CB1Rs on astrocytes, not on interneurons, are required for OD plasticity during the critical period.

**Figure 4.**
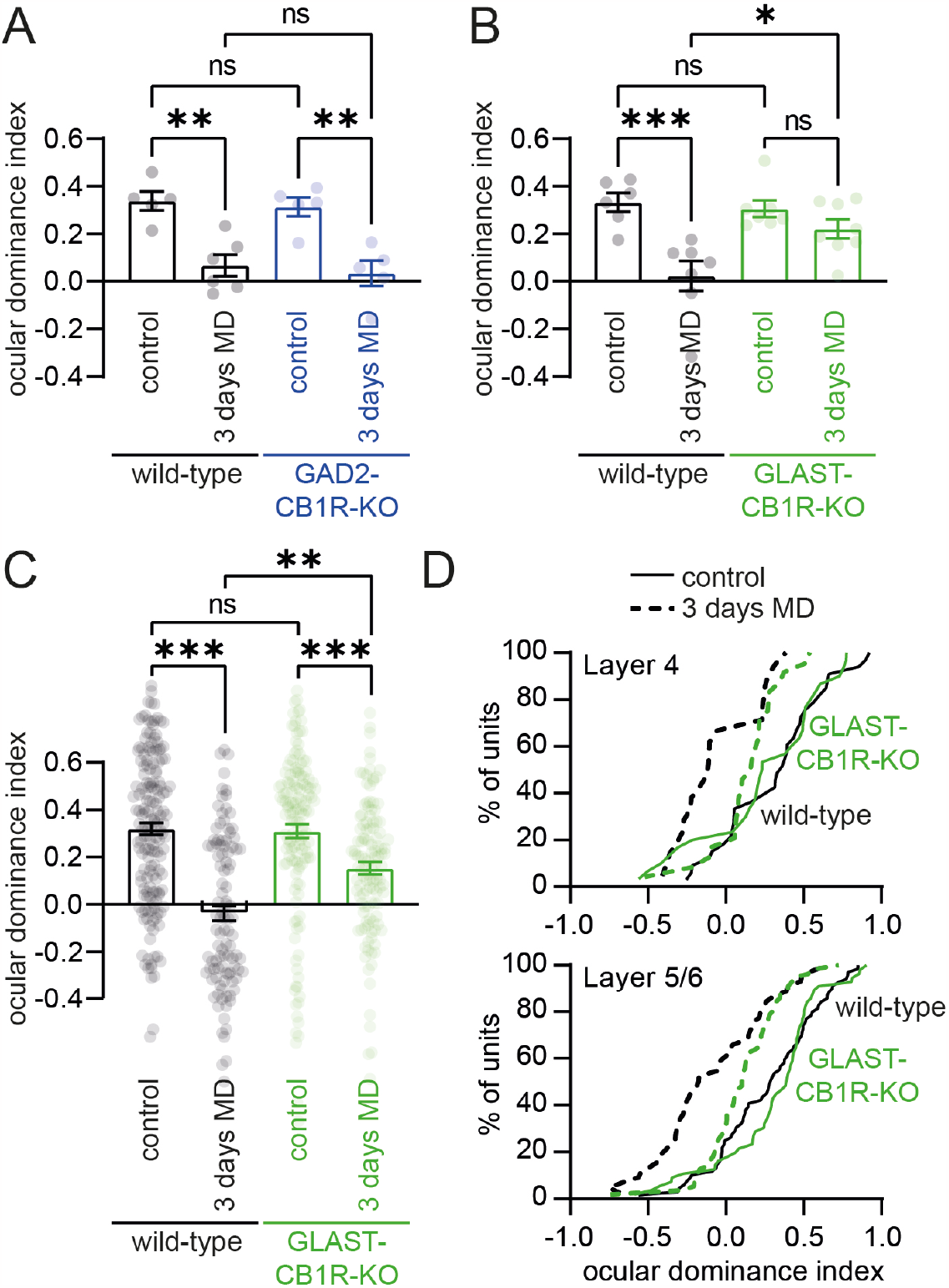
Ocular dominance plasticity is disrupted upon loss of astrocyte CB1 receptors. (A) Summary graphs of the ocular dominance index, as assessed using optical imaging of the intrinsic signal. Data is shown for GAD2-CB1R-KO mice (blue) and their wild-type littermates (black), both under control conditions and after three days of monocular deprivation (3 days MD). Dots indicate recorded ocular dominance index for individual mice, bars show means. (B) Same as in A, but now for GLAST-CB1R-KO mice (green) and their wild-type littermates (black). (C) Same as in A and B, but here ocular dominance index was assessed using *in vivo* electrophysiology. Dots represent individual single units, bars show means. (D) Cumulative distribution of ocular dominance index against % of units, for deep cortical layers (L4 and L5/6). Dotted lines represent undeprived controls, solid lines represent 3 days monocular deprivation. Green lines show data from GLAST-CB1R-KO mice, black lines from wild-type littermates.

### OD plasticity is disrupted in deep cortical layers upon loss of astrocytic CB1Rs

OD plasticity measured using optical imaging of intrinsic signal likely mainly reports plasticity in superficial cortical layers. Previous studies that described effects of pharmacological CB1R blockade on OD plasticity revealed that the effect of acute CB1R blockade on OD plasticity is layer specific, with OD plasticity in layer 2/3 of V1 being sensitive to treatment with a CB1R antagonist, while deeper layers show normal OD plasticity upon CB1R antagonist treatment^17,18^. To investigate whether the disruption of OD plasticity upon developmental loss of astrocytic CB1Rs was observed in deeper layers, we performed electrophysiological recordings using laminar probes in GLAST-CB1R-KO mice. Analyzing OD plasticity over all cortical layers confirmed the disruption of OD plasticity that we observed using intrinsic signal optical imaging. Upon removal of astrocyte CB1Rs, OD plasticity was still observed, but in significantly reduced form (wild-type: ODI non-deprived: 0.32±0.03, n=162/11; 3 days MD: −0.04±0.03, n=104/7; GLAST-CB1R-KO: non-deprived: 0.31±0.03, n=139/10; 3 days MD: 0.15±0.03, n=121/8; two-way ANOVA; interaction of genotype with OD shift *P*=0.002; Tukey’s post-hoc test: wild-type non-deprived vs 3 days MD: *P*<0.0001; GLAST-CB1R-KO non-deprived vs 3 days MD: *P*=0.0006; wild-type 3 days MD vs GLAST-CB1R-KO 3 days MD: *P*<0.0001; Fig. 4C). Next, we specifically looked at OD plasticity in deeper cortical layers, by separately analyzing units in layer 4 and layer 5/6, based on depth. We found that loss of astrocytic CB1Rs reduced OD plasticity in deep cortical layers (Fig. 4D; L4: wild-type: ODI non-deprived: 0.32±0.06, n=33/11; 3 days MD: −0.06±0.06, n=21/7; GLAST-CB1R-KO: non-deprived: 0.25±0.07, n=30/10; 3 days MD: 0.14±0.05, n=24/8; two-way ANOVA; interaction of genotype with OD shift *P*=0.033; L5/6: wild-type: ODI non-deprived: 0.28±0.04, n=69/11; 3 days MD: −0.09±0.05, n=44/7; -CB1R-KO: non-deprived: 0.31±0.04, n=56/10; 3 days MD: 0.11±0.03, n=56/8; two-way ANOVA; interaction of genotype with OD shift *P*=0.048). Therefore, genetic removal of astrocytic CB1Rs during development has a different and broader effect on OD plasticity than acute pharmacological CB1R blockade.

### Discussion

In this study we show that astrocytic CB1Rs contribute to the maturation of inhibitory synapses and affect OD plasticity in the developing V1. It is well known that the maturation of inhibitory synapses in V1 is required for the onset of the critical period of OD plasticity^1,19-22^. Our finding thus supports the idea that the deficit in OD plasticity observed in astrocytic CB1R deficient mice is caused by deficient maturation of inhibitory synapses affecting critical period onset.

During maturation of V1, inhibitory innervation changes extensively. During the first week after eye opening (p14-p21) the number and size of inhibitory synapses increase resulting in stronger inhibition^22^. The causal link between inhibitory maturation and opening of the critical period of OD plasticity is well established: In mice with reduced GABA-release due to the absence of GAD65, a protein involved in GABA synthesis, the critical period does not start^19,20^. Increasing GABAergic transmission in these mice by intraventricular benzodiazepine infusion rescues the phenotype and initiates the critical period^19^. In addition to the general increase in synaptic strength, synaptic release and short-term depression decrease with development, resulting in more reliable and precise inhibition^23-25^. At inhibitory synapses formed by parvalbumin-expressing fast-spiking interneurons the maturation of inhibitory synaptic strength and dynamics depends on CB1R signaling, since both processes are blocked by CB1R antagonist treatment or by genetic CB1R knockout^4,5,10^. This has been puzzling, since parvalbumin-expressing interneurons express no or only low levels of CB1Rs^11-13^. Our finding that inhibitory synaptic transmission is intact upon interneuron CB1R removal, but that it is disrupted when astrocyte CB1Rs are removed, provides an explanation for this apparent discrepancy.

The mechanism underlying CB1R-mediated inhibitory synapse maturation is not fully understood. In previous studies, CB1R inactivation resulted in both a blockade of iLTD and in a disruption of inhibitory synapse maturation. Therefore these processes were considered to be causally related^4,5^. Our observation that in GLAST-CB1R-KO mice inhibitory synapse maturation is disrupted, while iLTD is still intact, is surprising. Our results suggests that iLTD and inhibitory synaptic maturation are independent processes. Another possible explanation is that recombination in GLAST-CB1R-KO mice is incomplete, leading to residual CB1R expression in some astrocytes. Experiments with TdTomato reporter mice (Figure 1) reveal recombination in ∼80% of astrocytes upon a single early postnatal tamoxifen injection. Since full genetic CB1R removal from a cell requires bi-allelic recombination, while reporter gene expression is already induced by mono-allelic recombination, it is expected that ∼60-70% of astrocytes in GLAST-CB1R-KO mice fully lack CB1Rs. iLTD induced by strong electrophysiological stimulation of a large number of inhibitory synapses might be resilient to such incomplete CB1R inactivation, with CB1R activation on the remaining astrocytes being sufficient for iLTD induction.

Previous work has shown that acute pharmacological blockade of CB1Rs during the critical period reduces OD plasticity. In those experiments, OD plasticity was only affected in the superficial layers, while an OD shift could still be induced in layers 4-6^17,18^. LTD of excitatory synapses in layer 2/3 of sensory cortex is dependent on CB1Rs, while excitatory LTD (eLTD) in deeper layers is CB1R independent^26,27^. It is therefore believed that the effect of acute CB1R blockage on OD plasticity is caused by selective interference with eLTD in layer 2/3. We find that in astrocytic CB1R-deficient animals, OD plasticity is reduced in all layers of V1. This suggests that CB1Rs regulate OD plasticity through multiple mechanisms: acutely by mediating LTD at excitatory synapses in layer 2/3 neurons and indirectly by driving the development of inhibitory synapses in all cortical layers^4^. We do not know whether inactivating CB1Rs in astrocytes will also affect OD plasticity by an acute effect on excitatory LTD. But several studies have shown that astrocyte CB1Rs can regulate plasticity at excitatory synapses^7^, for instance by driving D-serine release necessary for NMDA receptor activation^28^. One would need to inactivate CB1Rs in astrocytes at a later age to dissociate developmental and acute astrocyte CB1R effects.

In GLAST-CB1R-KO mice, astrocyte CB1Rs are not only inactivated in V1, but also in the rest of the brain. We can therefore not rule out that the observed reduction of OD plasticity is caused by the absence of astrocytic CB1Rs in other brain structures, such as the dorsal lateral geniculate nucleus (dLGN) providing input to V1. We have shown that OD plasticity also occurs in the thalamus, and that thalamic synaptic inhibition is essential for OD plasticity both in dLGN and in V1^29^. However, in the absence of thalamic inhibition, OD plasticity was predominantly affected after 7 days of MD, while after 3 days of MD, the OD shift was barely reduced^29^. In our current study, we saw a strong decrease of OD plasticity already after 3 days of MD in GLAST-CB1R mice, suggesting that reduced thalamic inhibition is not the main cause of this plasticity deficit.

Taken together, we show that astrocytic CB1Rs are crucial for critical period plasticity in V1. These findings add to a larger body of research that reveal a role for astrocytes in regulating critical periods in the brain^30,31^. Interestingly, transplanting astrocytes from kittens into V1 of adult cats reopens the critical period of OD plasticity^32^. A more recent study found that transplanting immature astrocytes in V1 of adult mice reopens the critical period through degradation of the extracellular matrix^31^. Adult astrocytes no longer have the ability to re-open the critical period due to decreased metalloprotease 9 activity. Whether the expression of CB1Rs on immature astrocytes also contributes to their ability to alter critical period plasticity remains unclear. Future experiments involving the transplantation of astrocytes from young mice lacking CB1Rs may provide an answer.

## Methods

### Animals

Experimental procedures involving mice were in strict compliance with animal welfare policies of the Dutch government and were approved by the Institutional Animal Care and Use Committee of the Netherlands Institute for Neuroscience. Genetically-modified mice were bred on a C57Bl6/J background. For generation of conditional CB1R knockout mice, mice homozygous for a loxP-site flanked CB1R gene (*Cnr1*^flox/flox^ mice)^14^ and heterozygous for GAD2-Cre (Gad2-IRES-Cre mice; Jax Stock No: 019022)^15^ or transgenic for GLAST-CreERT2 (Jax Stock No: 012586) were bred. To assess efficacy and specificity of recombination in GLAST-CreERT2 mice these were crossed with ROSA-TdTomato reporter mice^33^, in which a Cre-dependent transgene encoding the tdTomato fluorescent protein is inserted in the ROSA26 locus (Jax Stock No: 007908). Experiments were performed on mice of either sex. Animals were housed on a 12 h light/dark cycle with unlimited access to standard lab chow and water.

### Tamoxifen injection

Mouse pups received a single intraperitoneal (i.p.) tamoxifen injection to induce Cre mediated recombination in the GLAST-CreERT2 line. Tamoxifen was dissolved in corn oil at a final concentration of 5 mg/ml. Dissolving tamoxifen was aided by placing the Eppendorf tube in an ultrasonic water bath, heated to 30°C, for ∼1 hour. Tamoxifen solution (20-25 μl) was injected using a thin insulin needle. To assess specificity and efficacy of recombination injections were performed either at P1 or between P3-P5. For all further experiments a single injection between P3-P5 was used.

### Monocular deprivation

Surgery for monocular deprivation (MD) was performed as follows: Mice were anesthetized using isoflurane (5% induction, 1.5–2% maintenance in 0.7 l/min O2). Edges of the upper and lower eyelids of the right eye (contralateral to the side on which recordings were performed) were carefully removed. Antibiotic ointment (Cavasan) was applied. Eyelids were sutured together with 2–3 sutures using 7.0 Ethilon thread during isoflurane anesthesia. Postoperative lidocaine ointment was applied to the closed eyelid. Eyes were checked for infection or opacity once reopened 3 days later, and only mice with clear corneas were included.

### Intrinsic signal optical imaging and electrophysiology

Mice were anesthetized by intraperitoneal injection of urethane (20% solution in saline, 1.2 g/kg body weight), supplemented by a subcutaneous injection of chlorprothixene (2.0 mg/ml in saline, 8 mg/kg body weight). Sometimes a supplement of about 10% of the original dose of urethane was necessary. Injection of anesthetic was immediately followed by a subcutaneous injection of atropine sulfate (50 μg/ml in saline, 1 μg/10g body weight) to reduce secretions from mucous membranes and facilitate breathing. Anesthesia reached sufficient depth after 45–60 min. Body temperature was monitored with a rectal probe and maintained at a temperature at 36.5 °C using a heating pad. The animal was fixated by ear bars with conical tips prepared with lidocaine ointment. A bite rod was positioned behind the front teeth, 4 mm lower than the ear bars. A continuous flow of oxygen was provided close to the nose. For analgesia of the scalp, xylocaine ointment (lidocaine HCl) was applied before resection of a part of the scalp to expose the skull.

OD measurements were performed as previously described^16^. In brief, the exposed skull was illuminated with 700 ± 30 nm light and the intensity of reflected light was measured. Responses were acquired with an Imager 3001 system (Optical Imaging, Israel). A gamma corrected computer screen was placed in front of the mouse, covering an area of the mouse visual field ranging from − 15 to 75 degrees horizontally and − 45 to 45 degrees vertically. First the retinotopic representation of V1 was mapped. Full contrast, square wave gratings of 0.05 cycles per degree (cpd), moving at 2 Hz and changing direction every 0.6 s were shown every 9 s for 3 s in a pseudo-randomly chosen quadrant while the rest of the screen was a constant gray. Fifteen stimuli in each quadrant were sufficient to construct a robust retinotopic map of V1. To subsequently measure OD, shutters were placed in front of both eyes. Either shutter opened independently at preset intervals, for a period of 6 seconds. After full opening of the shutter, the visual stimuli described above were presented in the upper nasal quadrant of the screen for a period of 3 seconds. Fifty responses to stimulation were recorded for each eye. For quantification, the response of a defined region of interest within the binocular part of V1 (as determined by retinotopic mapping) was normalized to the response seen in a region of reference (ROR) outside of V1, which lacked a stimulus specific response. The negative ratio of ROI over ROR signal was taken, normalized to the stimulus onset and averaged from the first frame after stimulus onset until 2s after stimulus offset. The Ocular Dominance Index (ODI) was calculated as (contralateral response − ipsilateral response)/(contralateral response + ipsilateral response).

For *in vivo* electrophysiology, a craniotomy was prepared over V1, 2.95 mm lateral and 0.45 mm anterior to lambda. Mice were placed in front of a gamma-corrected projector (PLUS U2-X1130 DLP), which projected visual stimuli onto a back-projection screen (Macada Innovision, the Netherlands; 60X42 cm area) positioned 17.5 cm in front of the mouse. One eye was covered with a double layer of black fabric and black tape while neuronal responses to stimulation of the other eye were recorded. Visual stimuli were created with the MATLAB (MathWorks) software package Psychophysics Toolbox 3^34^. The position of the receptive field was determined by displaying white squares (5 degrees) at random locations on a black background. The ODI was calculated by presenting each eye with alternating white and gray full-screen stimuli. Each stimulation lasted 3 seconds. There were 100 repetitions of both white and gray screens. Extracellular recordings from V1 were made using a linear silicon microelectrode (A1×16-5mm-25-177-A16, 16 channels spaced 50 m apart, Neuronexus, USA). Extracellular signals were amplified and bandpass filtered at 500 Hz-10 kHz before being digitized at 24 kHz using a RX5 Pentusa base station (Tucker-Davis Technologies, USA). A voltage thresholder at 3x standard deviation was used to detect spikes online. Custom-written MATLAB programs (http://github.com/heimel/inVivoTools) were used to analyze the data. We computed the spike - triggered average of the random spare square stimulus for receptive field mapping. The actual position and size of the visual field were calculated and corrected for the distance between the stimulus and the animal. We used the last 500ms of the previous trail as the baseline for each 3 s stimulus-related activity. As a result, we characterized visual responses as the difference between the first 500ms of the stimulus and the mean of the prior stimulus’s last 500ms activities. The greatest firing rates in the first 300ms of visual related reactions were regarded the peak visual responses to stimuli. The visual responses were calculated as 300ms average multi-unit responses. ODI was calculated as (*R*_*contra*_− *R*_*ipsi*_)/(*R*_*contra*_+ *R*_*ipsi*_), where the *R*_*contra*_ is the average multi-unit firing rate of the unit when contralateral eye was open and ipsilateral eye was covered; *R*_*ipsi*_ is opposite.

### Slice electrophysiology

For acute brain slice preparation, animals aged between P14-35 were briefly anesthetized using isoflurane, followed by decapitation. Brains were removed and placed in ice-cold slicing medium. For most experiments sucrose based slicing medium was used, containing (in mM): 212.7 sucrose, 26 NaHCO3, 3 KCl, 1.25 NaH2PO4, 1 CaCl2, 3 MgCl2 and 10 D(+)-glucose (carbogenated with 5% CO2/95% O2; osmolarity 300-310 mOsm). For some experiments choline chloride based slicing medium, containing (in mM): 110 choline chloride, 7 MgCl2, 0.5 CaCl2, 2.5 KCl, 11.6 Na-ascorbate, 3.10 Na-pyruvate, 1.25 NaH2PO4, 25 D-glucose and 25 NaHCO3 (carbogenated with 5% CO2/95% O2; osmolarity 300-310 mOsm) was used. Quality of slices and properties of recorded neurons were indistinguishable for both solutions. Coronal slices (350 μm) containing V1 were prepared using a vibratome. Within each slice, the hemispheres were separated with a scalpel at the middle axis to be used for individual recordings. The slices were transferred to a holding chamber and left to recover for at least 30 min at 35°C in carbogenated artificial cerebrospinal fluid (ACSF), containing (mM): 124 NaCl, 26 NaHCO3, 3 KCl, 1.25 NaH2PO4, 2 CaCl2, 1 MgCl2 and 10 D(+)-glucose (carbogenated with 5% CO2/95% O2; osmolarity 300-310 mOsm). After recovery the holding chamber was moved to room temperature and slices were kept until recording (up to 8 hours after slice preparation).

For recording, slices were transferred to the stage of an upright microscope, where they were continuously perfused with heated (30-32°C) ACSF. NMDA receptor and AMPA receptor mediated glutamatergic synaptic responses were blocked by addition of D-AP5 (50 μM) and DNQX (10 μM) to the recording ACSF. Whole cell patch-clamp recordings from pyramidal neurons in Layer 2/3 of V1 were made using a Multiclamp 700B amplifier in voltage clamp mode and PClamp software (Molecular Devices, USA). Cells were patched with borosilicate glass electrodes with tip resistances of ∼3.5 MOhm, and filled with intracellular solution containing (mM): 120 CsCl, 8 NaCl, 10 HEPES, 2 EGTA, 10 Na-phosphocreatine, 4 Mg-ATP, 0.5 Na-GTP and 5 QX-314 (pH: 7.4; osmolarity ∼285 mOsm). IPSCs were evoked with an ACSF filled glass electrode with a broken tip or with a concentric bipolar stimulation electrode, placed in Layer 4. Intensity of the stimulation pulse was adjusted to obtain a reliable and stable IPSC response.

For experiments in which iLTD was evoked, IPSCs were evoked every 20 seconds until a stable baseline was established. Next, iLTD was induced using theta-burst stimulation (TBS), consisting of 8-10 thetaburst epochs delivered every 5 seconds. Each theta-burst epoch consisted of 10 trains of 5 pulses at 100 Hz, with the trains being delivered at 5 Hz (adapted from^4,5^). In a small subset of experiments, the strength of extracellular stimulation was doubled during TBS delivery. Because this did not affect the magnitude or dynamics of iLTD, all experiments were grouped for final analysis. AM251 was diluted in DMSO, and the stock solution was added to ACSF to obtain a final concentration of 10 μM (final DMSO concentration: 0.1%).

IPSC amplitude was analysed using custom scripts in IGOR pro (WaveMetrics, USA). For iLTD analysis, magnitude of iLTD was determined by comparing the IPSC amplitude during the 10 minute baseline to the IPSC amplitude 10-20 minutes after TBS. Recordings were excluded if the IPSC amplitude differed >12.5% between the first and the last 6 responses of the baseline, if input resistance increased >25% between baseline and iLTD window, if access resistance increased to >20 MOhm, or if leak current reached lower than −500 pA.

### Immunohistochemistry

Mice were anesthetized with an overdose of pentobarbital (100 mg/kg i.p.), followed by transcardial perfusion with 4% paraformaldehyde (PFA) in phosphate buffered saline (PBS). Brains were isolated and post-fixated for >2 hours in PFA at 4°C. After changing to PBS, coronal slices (50 μm thickness) were prepared. Slices were incubated for 2 hours in 500 μl blocking solution (0.1% Triton X-100, 5% NGS in PBS) on a rotary shaker at room temperature. Afterwards, slices were incubated with primary antibody containing solution and left overnight at 4°. The next day primary antibody solution was discarded, and slices were washed three times for 10 min at room temperature on the rotary shaker with 500 μl of washing solution (0.1% Tween in PBS). Secondary antibody solution (250 μl) was added per well and slices were incubated for 1 h at room temperature on the rotary shaker. Next, slices were again washed three times for 10 min at room temperature on the rotary shaker with washing solution. Stained slices were mounted on glass slides using Mowiol. The following antibodies were used: Glutamine Synthetase (monoclonal mouse, MAB302, Merck Millipore, USA), NeuN (monoclonal mouse, 1:1000, MAB377, Merck Millipore, USA), NG2 (polyclonal rabbit, 1:250, AB5320, Merck Millipore, USA) and Olig2 (monoclonal mouse, 1:250, MABN50, Merck Millipore, USA). Secondary antibody conjugated with Alexa-488 was used (1:250 or 1:500, ThermoFisher, USA). Imaging of the immunostained sections was done using a Leica TCS SP5 Confocal microscope (Leica, Germany).

### Statistics

Data representation and statistical analysis were performed using GraphPad Prism 9.0 (GraphPad Software, USA). Parametric or non-parametric statistics were used depending on whether data points were normally distributed. When necessary, correction for multiple comparisons was applied (Tukey’s post-hoc test for two-way ANOVA). The used tests are indicated in the text. Statistically significant differences were defined as P ≤ 0.05. Data are represented as mean ± SEM.

## Acknowledgements

The authors thank Giovanni Marsicano and Beat Lutz for providing *Cnr1*^flox/flox^ mice, and Emma Ruimschotel for excellent technical assistance. This study was funded by an NWO Veni grant to RM (863.12.006).

